# Distinctive functional regime of endogenous lncRNAs in dark regions of human genome

**DOI:** 10.1101/2020.12.06.413880

**Authors:** Anyou Wang, Rong Hai

## Abstract

Eukaryotic genomes gradually gain noncoding regions when advancing evolution and human genome actively transcribes >90% of its noncoding regions^1^, suggesting their criticality in evolutionary human genome. Yet <1% of them have been functionally characterized^2^, leaving most human genome in dark. Here we systematically decode endogenous lncRNAs located in unannotated regions of human genome and decipher a distinctive functional regime of lncRNAs hidden in massive RNAseq data. LncRNAs divergently distribute across chromosomes, independent of protein-coding regions. Their transcriptions barely initiate on promoters through polymerase II, but mostly on enhancers. Yet conventional enhancer activators(e.g. H3K4me1) only account for a small proportion of lncRNA activation, suggesting alternatively unknown mechanisms initiating the majority of lncRNAs. Meanwhile, lncRNA-self regulation also notably contributes to lncRNA activation. LncRNAs trans-regulate broad bioprocesses, including transcription and RNA processing, cell cycle, respiration, response to stress, chromatin organization, post-translational modification, and development. Overall lncRNAs govern their owned regime distinctive from protein’s.

## Introduction

Human cells consume enormous energy to transcribe over 93% of its genome that consists of 98% noncoding regions^1^, suggesting the noncoding region as the crucial component in human biology. Long coding RNAs (lncRNAs) predominate the transcripts from noncoding regions^3^. Understanding the abundant lncRNA functions helps to appreciate the fundamental basis of human genome.

The general strategy for characterizing protein-coding mRNA has been conventionally applied to understand lncRNAs^2,4^. For example, lncRNA identification have been derived from mRNA features, such as promoter, start coden, poly-A and RNA polymerase II (Pol II), and DNA conservation^2,4^. Combining mRNA concept and sequencing approaches, the latest GENCODE project V35^3^ has collected 16,899 lncRNAs, in which lincRNAs (long intergenic non-coding RNA) and antisense RNAs have been merged into a lncRNA category. FANTOM project has also identified 19,175 lncRNAs from 5’s strategy^5^. However, most lncRNAs originate from enhancers rather than from promoters^5^ and lncRNA could initiate from 3’s and other mechanisms. These current experimental approaches only identified a limited number of lncRNAs. Bioinformatics and computational tools can help to identify lncRNAs^6,7^, but their development has been slow to systematically identify novel lncRNAs in human genome, leaving most genome regions in dark.

On the other hand, most lncRNAs identified to date in humans are tissue-specific and are not conserved^8^. However, a certain number of lncRNAs are evolutionary or functionally conserved. For example, zebrafish carry conserved lncRNAs crucial for embryonic development and these lncRNAs are functionally conserved across species^2^. Mouse genome contains >1,000 lncRNAs with substantial evolutionary conservation (>95%) cross-mammalian^9^. Human genome contains so many noncoding regions that we hypothesize that it holds a large number of functional lncRNAs endogenous across all cell-types and tissues and conditions.

This study employed our new software FINET^10^ to systematically identify the functionally unannotated lncRNAs(ulncRNAs) endogenous in dark regions of human genome via exhaustively searching a ulncRNA regulatory network hidden in massive data, including all RNAseq data from SRA database. We then generated quantitative patterns from this network to uncover distinctive mechanisms of ulncRNA activation, regulation, and function.

## Results

### Endogenous ulncRNA regulatory network

The data from our previous study revealed that only 22% of lncRNAs annotated by GENCODE project are functionally endogenous(**Figure _S1**)^11^, indicating that most annotated lncRNAs are tissue-specific. The number of endogenous ulncRNAs in dark regions of human genome remains unknown.

To identify endogenous ulncRNAs, we developed an algorithm strategy to systematically capture all functional ulncRNAs endogenous across all human tissues and conditions **(Figure 1A).**This strategy includes the following 4 key steps. (1) split unannotated dark regions (100bp distance from annotated regions) into 300bp RNA fragments as preliminary ulncRNAs (**Figure 1A**, **materials and methods**). (2) identify interactions of ulncRNAs endogenous in human genome from massive data (all RNAseq data deposited in SRA) by using our FINET software that infers endogenous regulatory interactions from massive data with high accuracy^10^(materials and methods), and simultaneously remove unfunctional ulncRNAs with no regulatory interactions (e.g. removing the blue color ulncRNA in Figure 1A). (3) concatenate conjunctions(e.g. concatenate ulncRNA1 and ulncRNA2 into ulncRNA12 in **Figure 1A**). (4) assemble all interactions into a ulncRNA regulatory network, in which an individual ulncRNA possesses at least one functionally regulatory interaction. This strategy generated an endogenous ulncRNA regulatory network, which includes 16594 unique ulncRNAs with 62586 edges and 29794 nodes deposited in the project website^12^(**Figure 1A-1B**).

**Figure 1.**
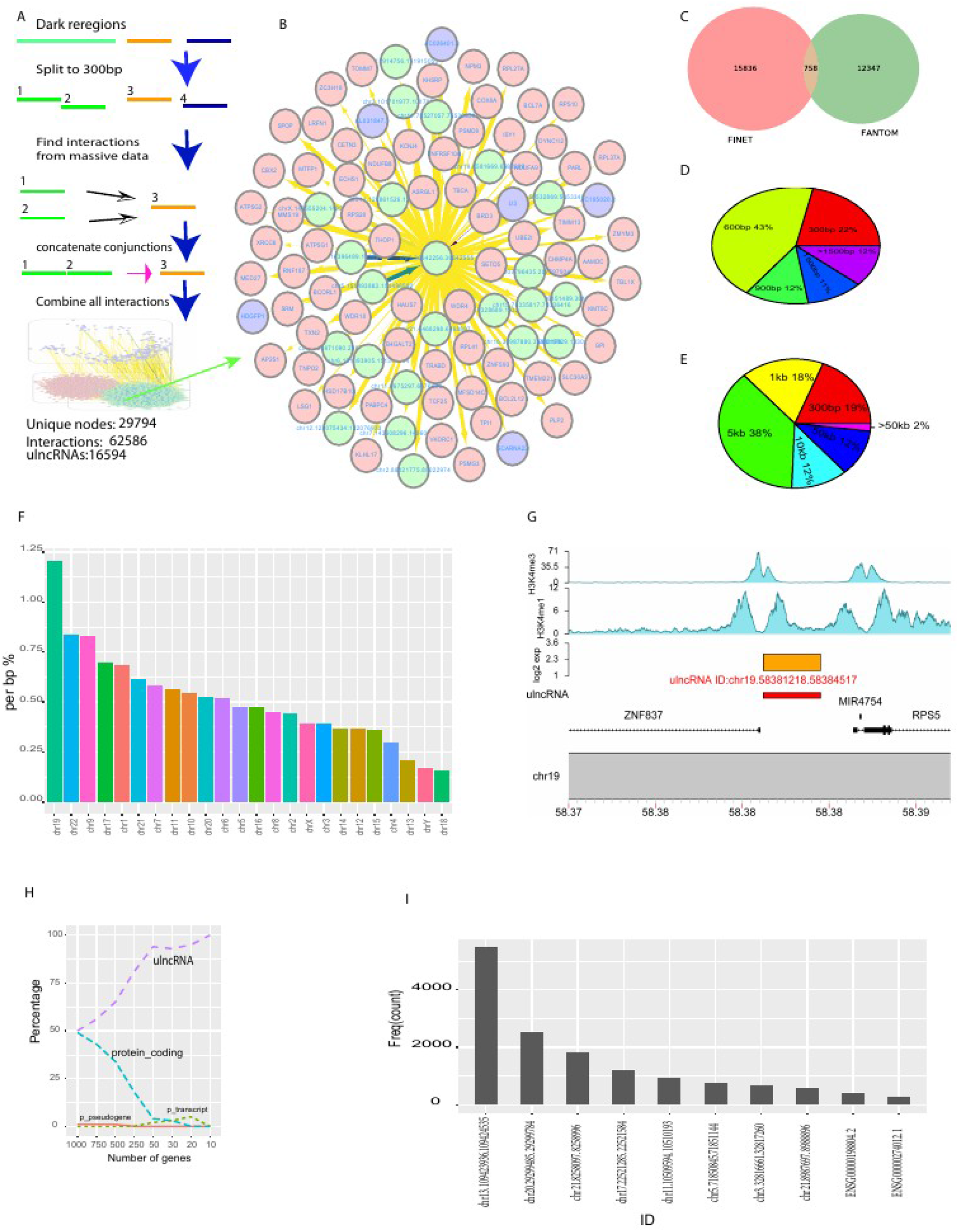
Functionally endogenous ulncRNAs identified in human genome. A, workflow of ulncRNA regulatory network identification. B, a sample network of an ulncRNA (ID: chr3.36642256.36642555). ulncRNA ID was named as chromosome plus coordination. Node color denotes gene category, pink:protein, green:ulncRNA, purple:annotated noncoding RNAs. Edge color represents significance (p-value), from low (yellow) to dark blue (high, low p-value). edge thickness denotes confidence measured by frequency score in our FINET software^10^, thicker, more confident. C, Overlap of lncRNAs from FANTOM and total 16594 unique ulncRNAs identified by this present study via using FINET software. D, size distribution of total 16594 ulncRNAs. E, Distribution of minimum distance from ulncRNAs to proteins. F, ulncRNA density along chromosomes measured by total length of RNAs/chromosome length (Figure S2 for detail). G, an example profiling of ulncRNA, chr19:58381218.58384517. Its log2 expression level was shown in brown, and the profiling of two typical histone markers was plotted, H3K4me1(ENCODE ID: ENCFF730CTY.bigWig) and H3K4me3 (ENCODE ID: ENCFF881MFX.bigWig). Annotated genes ZNF837, RIPS5, MIR4754 was marked in the bottom. H, gene category proportion of top 1000 centrality in ulncRNA network. P_ denotes processed. For example, p_pseudogene as processed_pseudogene. I, top 10 highest connected nodes in ulncRNA network. Frequency count represents interactions(degrees).

From the overall outlook of entire network, ulncRNAs were mostly clustered with proteins and ulncRNAs, indicating that ulncRNAs primarily regulated themselves and proteins(**Figure 1A bottom and Figure 1B**).

Among 16594 ulncRNAs, only 4.5% (758/16594) overlapped with lncRNAs identified by FANTOM project (**Figure 1C**), which used experiments to identify lncRNAs. This indicated that most of lncRNAs identified by both FANTOM and GENCODE project are cell type specific.

Overall, we revealed a novel ulncRNA regulatory network endogenous across all human genomes and conditions, with no cell type specificity.

### Key characteristics of endogenous ulncRNAs

To understand the primary characteristics of these 16594 ulncRNAs, we examined the distribution of their length, closest distance to protein-coding sequences, chromosome distribution, and key hubs in the network. Most of ulncRNAs (65%) were less than 600bp length (43% of 600bp + 22% 300bp) (**Figure 1D**), and long ulncRNAs (>1500bp) occupied 12%. Surprisingly, most of these ulncRNAs distribute far away from protein-coding regions, with >62% (62% =38%+12%+12%) located >5kb from protein-coding regions(**Figure 1E**). We also calculated the ulncRNA density across chromosome (total ulncRNAs length/chromosome length) and found that chr19 possessed the most density of ulncRNAs, with more than 1bp ulncRNAs in every 100bp DNA length (>1% per bp, **Figure 1F**, **Figure S2** for full chromosome distribution). An example in chr19 (id:chr19.58381218.58384517) was shown in **Figure 1G.** This highly expressed ulncRNA corresponded to H3K4me1 binding in the same location (**Figure 1G**).

To understand the crucial hubs in the ulncRNA network, we examined both the centrality of the entire network and the highest connected nodes. ulncRNAs worked as the key hubs in this network and they occupied more than 50% of top 1000 centrality and 90% of top 50 as ulncRNA centrality(**Figure 1H**, materials and methods). In addition, 8 out of top 10 highest connected nodes were ulncRNAs (**Figure 1I**). The top 1 of these ulncRNAs, chr13.109423936.109424535, connected with total 5511 components in the entire network, including 5319 individual proteins targeted by ulncRNAs. This indicated that ulncRNAs govern the ulncRNA regime, instead of proteins as thought.

Overall the ulncRNA regime is distinctive from proteins, in which ulncRNAs are short, far away from coding regions, varied in chromosome distribution, and controlled by ulncRNA themselves.

### Systematic mechanisms of ulncRNA transcription origination

The mechanisms of lncRNA origination have been intensely debated^2^. Enhancers and RNA polymerase II have been thought as the primary factors for lncRNA activation^2^. To understand the mechanisms of ulncRNA origination, we systematically examined the contribution of both epigenetic and genetic transcription factors to ulncRNA activation. A factor contribution was measured by its binding abundance within 1000bp from ulncRNA TSS (transcription start site). For unbiased results, we included as many as possible cell types and conditions and downloaded all 780 samples of top 9 factors measured most frequently by Chip-seq from ENCODE(**Table_s1**), which contained 4 markers for enhancer (ATAC_seq^13^, H3K4me1^14^, H3K27ac^15^, H3K9ac^16^), 3 for promoter (H3K4me3^17^, POLR2A(polymerase II), H3K36me3 (exon)^18^), and 2 for silence and tissue specificity (H3K27me3 ^19^ and H3K9me3^2,19^).

To better understand the activation mechanism of ulncRNAs, we compared lncRNA binding profiling to protein’s and used the same number of genes for both ulncRNAs and proteins. From our previous study, we learned that 14122 proteins were active in normal conditions^11^, thus we randomly selected 14122 ulncRNAs out of 16594 to match the protein number(materials and methods). For each marker, we counted its binding sites along these 14122 ulncRNAs/proteins (<=1000bp TSS) for each sample. A box-plot of these binding sites of these 9 markers for all samples showed that POLR2A barely bound to lncRNAs, with median at 10% of ulncRNAs (1479/14122, **Figure 2A**). P value 0.1 could be treated as outliers and 10% bindings indicated that polymerase II plays much less role than thought in activating ulncRNA transcription.

**Figure 2.**
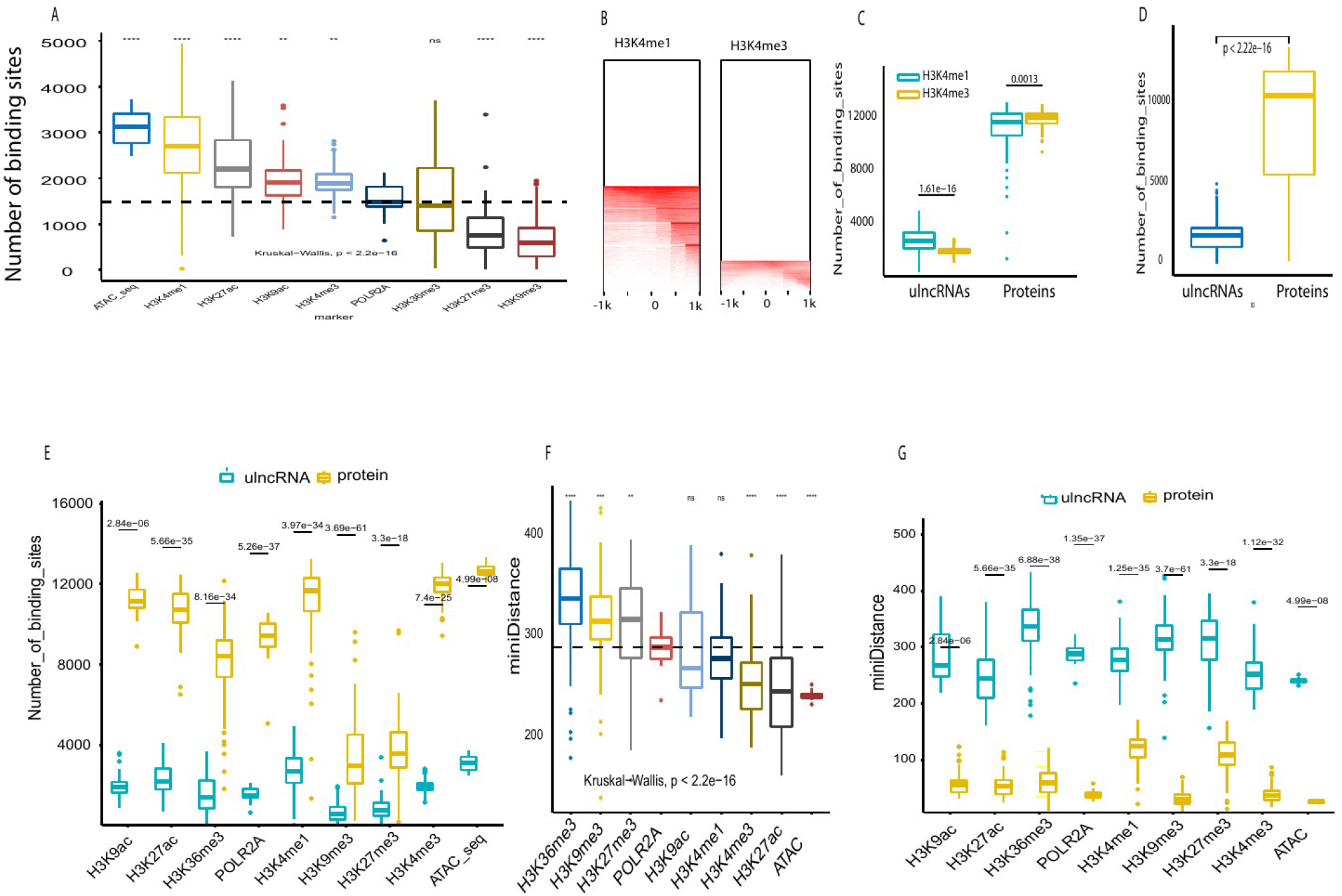
Contributions of histone and transcript factor to the ulncRNA activation. A, the frequency (total number of binding sites) of 9 markers that bind to ulncRNA promoter regions (within 1000bp from TSS, transcription start site). The black line represents the median of POLR2A binding sites(1479). These 9 markers were extracted from ENCODE chip-seq database (**table_S1**). Significance level, **,**** denotes pvalue <0.0034 and 3.4e-06 respectively. All labeled pvalue were derived from t test unless specific noted in this study. B, binding heatmap of H3K4me1 and H3K4me3. The heatmap was plotted by using the median of 1918 and 2694 samples respectively for H3K4me1 and H3K4me3 measured by ENCODE project (table_s1 H3K4me1bam and H3K4me3bam). C, Comparison H3K4me1 and H3K4me3 binding sites of ulncRNAs and proteins. Labeled number represents pvalue derived from t test in this study. D, Comparison of total binding sites of 9 markers between ulncRNAs and proteins. E, Binding comparison of each marker between ulncRNAs and proteins. F, minimum distance (bp) from marker binding to TSS. **,***,**** denotes p<0.0055, 0.00053, and 5.7e-08 respectively. G, minimum distance (bp) comparison of each marker between ulncRNAs and proteins.

Surprisingly, all three enhancer biomarkers, ATAC, H3K4me1, H3K27ac, H3K9ac, exhibited significant higher binding sites than POLR2A(**Figure 2A**). Moreover, ATAC and H3K4me1 (a well recognized marker for enhancers) had the highest bindings, with median binding rate of 22%(3119/14122) and 19% (2693/14122) respectively(**Figure 2A**). Furthermore, H3K4m1 bindings was significantly higher than H3K3me3, a marker for active promoters near TSS, (**Figure 2B-2C**, **Figure 1G**), while H3K4me3 bindings was higher in proteins (**Figure 2C**). This indicated that enhancers play a much greater role than polymerase II in activation of ulncRNAs.

To better understand the whole picture of these factor bindings, we box-plotted all the binding sites for ulncRNAs and proteins. ulncRNAs contained significantly lower bindings than proteins (**Figure 2D-2E**), at median of 1757 and 10321 for ulncRNAs and proteins respectively(**Figure 2D**), indicating that only 12% (1757/14122) of ulncRNAs possessed one biomarker to bind while 73% (10321/14122) of proteins have at least one biomarker to bind. Actually, all these histone marker bindings to ulncRNAs were significantly lower than to proteins (**Figure 2E**, **Figure S3**). For example, H3K4me3, H3K4me1 and POLR2A densely bound to proteins with 85%(12035/14122), 83%(11666/14122) and 67% (9446/14122) respectively(**Figure S3**). Moreover, proteins simultaneously possessed multiple factors for enhancers (e.g. ACTA and H3K4me1) and promoters (H3K4me3) to densely bind (**Figure 2E**), but ulncRNAs possessed much less factors to bind. This might partially interpret the low expression level of ulncRNAs. However, the overall low bindings (12%) and the 19% of H3K4m1 bindings were obviously not sufficient to activate the widespread ulncRNAs, suggesting that the key mechanism accounting for the majority of ulncRNAs activation remains to be investigated.

To appreciate the binding distance to ulncRNA TSS, we calculated the minimum distance from factor bindings to TSS. The minimum distance median ranged from 240pb (ATAC) to 336bp (H3K36me3) (**Figure 2F**). Factors with most binding sites(**Figure 2A**), including ATAC, H3K4me1, H3K27ac, H3K9ac, and H3K4me3, bound to ulncRNAs with short distance to TSS(**Figure 2F**). However, these short distances for ulncRNAs were significantly longer than proteins, in which factors bound much closer to protein TSS (**Figure 2G**, **Figure S4**, **Figure S5**). Furthermore, four markers (ATAC, H3K9me3, H3K4me3, PLOR2A) bound to protein TSS within 50bp (median) while the rest within 120bp (median) (**Figure S4**). In contrast, the median for all ulncRNAs was close to 280bp (**Figure S5**). This was another line of evidence for ulncRNA initiation regions distinctive from proteins. Altogether, these above suggested that ulncRNA activation mechanism is different from proteins and that the alternative mechanism for ulncRNA activation remains dark.

### ulncRNA regulators

To understand the regulators for ulncRNAs, we examined all ulncRNA regulators in the entire ulncRNA regulatory network and found a total of 31051 ulncRNA regulators. The most abundant regulators (>31%) were located outside their owned chromosomes(**Figure 3A**), suggesting that regulators trans-regulate ulncRNAs. Among 31051 regulators, 65% of them are ulncRNAs(**Figure 3B**), suggesting ulncRNAs primarily regulate themselves, consistent with our previous studies showing that noncoding genes tends to trans-regulate themselves at the same category^11^.

**Figure 3.**
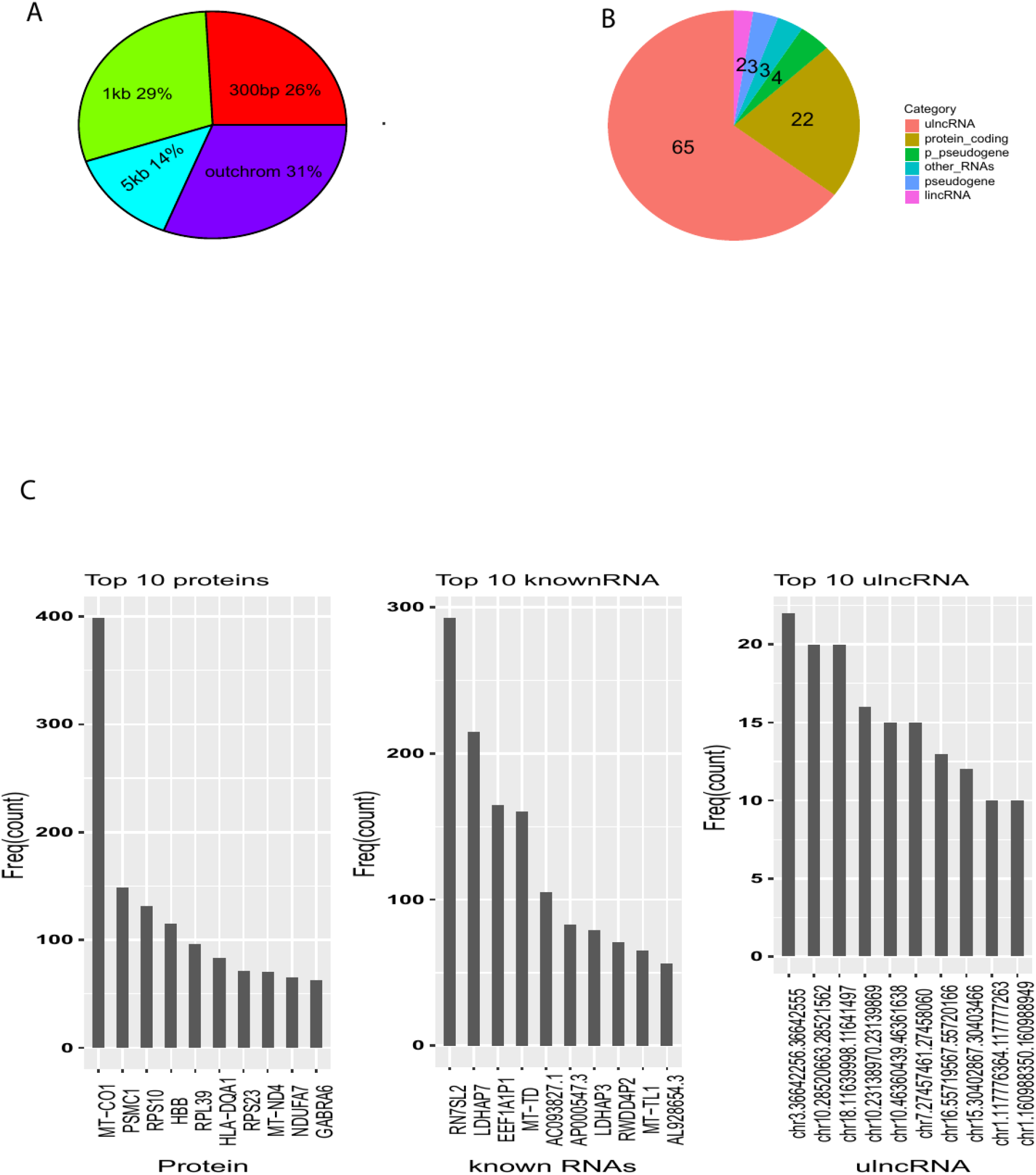
ulncRNA Regulators. A, distribution of distance from ulncRNA regulators to ulncRNAs (percentage of total 31051 ulncRNA regulators). B, gene categories of ulncRNA regulators (% of total 31051 ulncRNA regulators). C, top 10 highest connected regulators of proteins, annotated RNAs, and ulncRNAs.

In contrast to the conventional notion that proteins serve as primary regulators for lncRNAs, proteins actually work as secondary regulators (22%) for ulncRNAs (**Figure 3B**). Among protein regulators, a mitochondrial protein MT-CO1 connected to most ulncRNAs, with 400 interactions (**Figure 3C left panel**). Another mitochondrial protein MT-ND4 was also ranked as top 8 highest connected regulators (**Figure 3C left panel**). Moreover, two annotated noncoding RNAs, MT-TD and MT-TL1, were also in top 10 noncoding regulators for ulncRNAs(**Figure 3C right**). This suggested that mitochondrial components play a critical role in regulating ulncRNAs.

### ulncRNA targets

Whether lincRNA target their neighbor genes is debated ^2,8^. We plotted gene expression regression of ulncRNAs and their closest proteins within distance of 300bp and 1000bp respectively, and found that ulncRNAs did not regulate their neighbor proteins (**Figure 4A**, **Figure S6**). Instead ulncRNAs regulated their targets in a trans-regulatory manner, with the majority of ulncRNAs (57%) across chromosomes(**Figure 4B**). This parallels with a recent observation showing that majority of lncRNAs locate in cytoplasm as trans-regulators^20^. This is also consistent with our study on annotated lncRNA trans-regulation mechanism^11^.

**Figure 4.**
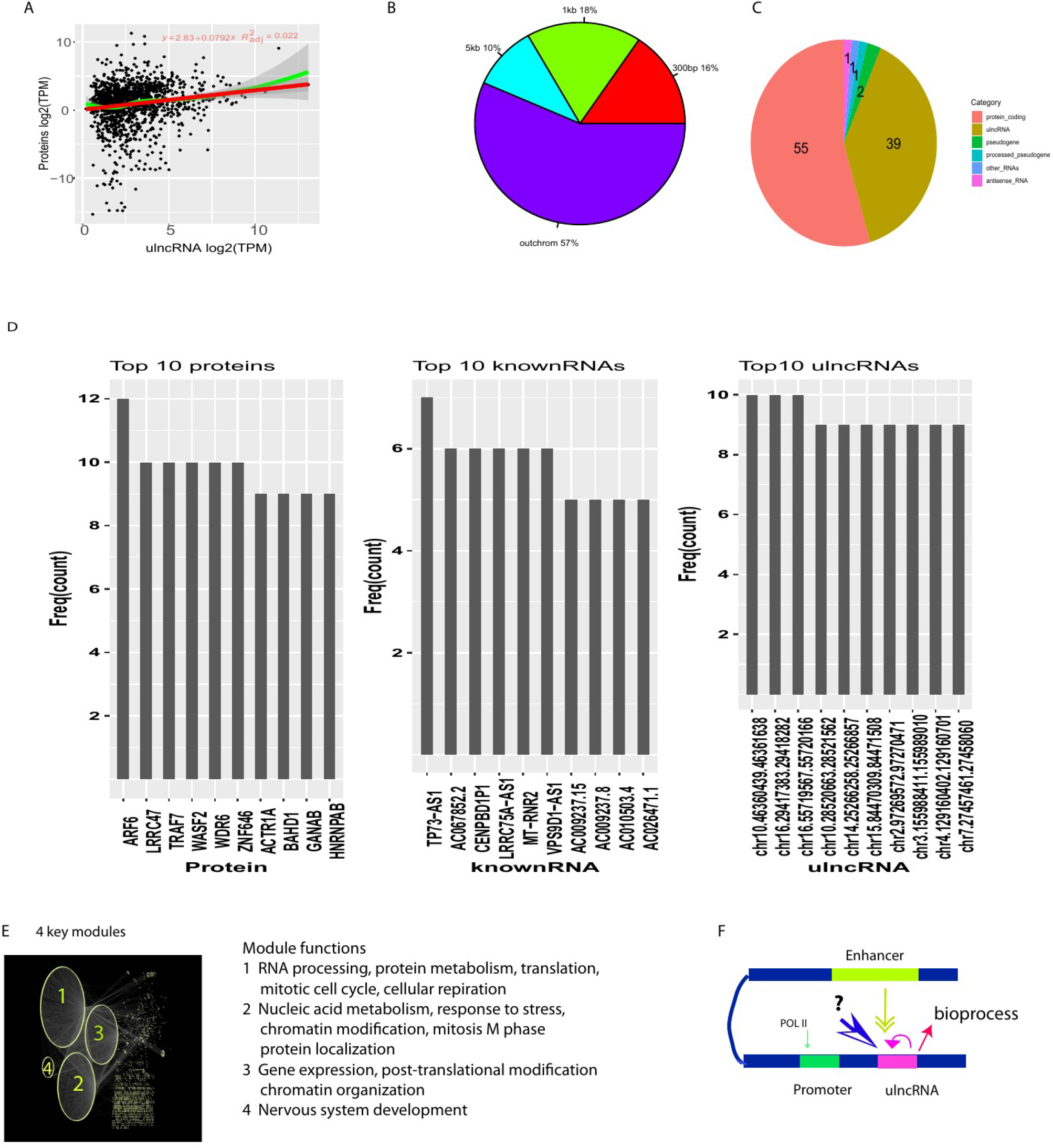
ulncRNA targets. A, Gene expression correlation between ulncRNAs and their closest proteins (within 300bp). B, Distribution of distances from ulncRNAs to their targets (percentage of total 51633 ulncRNA targets). C, gene categories of ulncRNA targets (% of total 51633 ulncRNA targets). D, top 10 highest connected ulncRNA targets of proteins, annotated RNAs, and ulncRNAs. E, top 4 functional modules in ulncRNA network. The size of module represents its member abundance. F, Functionally scheme of ulncRNA. The arrow size and line thickness represent the quantitative weight of importance. For ulncRNA activation, the factor importance ranking is following, unknown factor > ulncRNA > histone > POL II.

The majority of ulncRNAs target proteins (55%, **Figure 4C**), but their targets are scattered, with only 12 interactions of top 1 protein targets and less than 10 interactions with annotated noncoding RNAs and ulncRNAs (**Figure 4C**), suggesting that ulncRNAs primarily perform broad-but fine-regulation toward their targets.

### ulncRNA primary functions

To understand the primary functions of ulncRNAs, we investigated the key biological functions of ulncRNA targets. Among ulncRNA targets, proteins dominated the whole profiling (>55%, **Figure 4C**) and protein functions have well been characterized. These ulncRNA-targeted protein functions should represent the primary functions of ulncRNA targets. We searched ulncRNA target modules by spectral partitioning algorithm^21,22^ and found 4 key modules with biological functions (**Figure 4D**, **Figure S7-S10**). Their functions were primarily relevant to RNA processes but included 7 key categories,1) transcription and RNA processing (RNA splicing, ncRNA metabolic and processing); 2) mitotic cell cycle and DNA replication; 3) cellular respiration; 4)cellular response to stress (DNA repair); 5)chromatin organization; 6) translation and post-translational protein modification, proteasomal protein catabolic process, and protein localization; 7)nervous system development. These uNRA target functions suggested that biological roles of ulncRNAs are broader than thought.

To summarize, histone modifications on enhancers play more important role in activating ulncRNAs than polymerase II in promoters, but these histone modifications only account for a small proportion (<20%) of ulncRNA originations. The ulncRNA-self regulation and unknown mechanism contribute most of ulncRNAs activation and an array of biological functions(**Figure 4E**).

## DISCUSSION

This present study systematically decoded functionally endogenous lncRNAs from unannotated regions of human genome and revealed a functionally distinctive regime for lncRNAs. LncRNAs are widely expressed along the human genome but only a small proportion of lncRNAs have been identified and these annotated lncRNAs are mostly tissue specific^4,5^. Little has been known about lncRNA endogenicity in human genome. This present study overcame the limitations of tissues and conditions by using all RNAseq data from SRA and revealed 16594 endogenous lncRNAs in human genome. These lncRNAs mostly self-regulate independently of proteins and establish their own functional regime distinctively from protein’s in terms of distribution, activation, regulation and function.

The mRNA origination concept has been widely applied to lncRNA study^2^. Several mechanisms have been proposed for lncRNA origination, such as promoter and POL II and protein-based histone modifications on enhancers^2,4^. However, our data revealed that these conventional mRNA-based mechanisms only account for around 20% of lncRNA activation. The primary mechanism remains to be investigated, but lncRNA self-regulation may contribute to their activation. Individual lncRNAs are heavily regulated by other individual ulncRNAs. The lncRNA expression levels primarily result from lncRNA-self regulation. In normal conditions, these regulations stay weak to save energy, but under stimulation a certain group of lncRNAs would be highly activated by an array of lncRNA individuals and perform their biology functions.

lncRNAs are expressed at much lower levels than protein mRNAs. The mechanism for that remains debated^2^. The short life-span and the low promoter transcription efficiency have been thought as the mechanism of low lncRNA expression, but recent studies have demonstrated that lncRNAs have a similar half-life as normal mRNA and the bi-directional promoter working for proteins perform similarly for lncRNAs^2^. These two mechanisms hardly interpret the low lncRNA expression. The critical mistakes in these two mechanisms resulted from an assumption that lncRNAs employ the same promoter as proteins do. Our data showed that ulncRNAs rarely use protein promoters but they primarily employ enhancers far away from normal protein promoters. This parallels with recent observations showing lncRNA initiations from enhancers^5^. However, this enhancer origination for lncRNAs is distinctive from protein. In protein regime, histone modification like H3K4me2 are sufficient for transcription initiation, but all these protein-based factor bindings to lncRNAs are too low to initiate widespread lncRNAs regardless of enhancers and promoters. These low bindings of all transcription factors at least interpret partial mechanism of low lncRNA expression.

lncRNAs were once thought of as junk with no functions, but recently their functions have been recognized as regulators in several important processes such as growth and metabolism^23–26^. Our recent study also unearthed noncoding RNAs as the universal deadliest regulators for all cancers^27^. However, the primary functions of the vast majority of human genome occupied by lncRNAs still remain unknown. Here, we systematically reveal they target proteins functioning in an array of bioprocesses, such as transcription and RNA processing, mitotic cell cycle and DNA replication. These help us to understand the basic mechanism of lncRNA biological functions.

LncRNAs pre-dominate most of the human genome and have their own regime distinct from proteins. Applying protein strategy and concept to understand lncRNAs may be misleading. We need create a novel concept system to understand lncRNAs and human functional genome.

## Data and material availability

Data deposited in the project website^12^

Materials and methods in the project website^12^ and supplemental section

## Supplemental figure legends

**Figure_s1, overlap of annotated endogenous lncRNAs identified by FINET and total lncRNA annotated by GENCODE V27**

## References

1. 3 Characterization of intergenic regions and gene definition. Nature 1–1 (2019) doi:10.1038/nature28172.

2. Ransohoff, J. D., Wei, Y. & Khavari, P. A. The functions and unique features of long intergenic non-coding RNA. Nat. Rev. Mol. Cell Biol. 19, 143–157 (2018).

3. GENCODE - Human Release 35. https://www.gencodegenes.org/human/.

4. Schlackow, M. et al. Distinctive Patterns of Transcription and RNA Processing for Human lincRNAs. Mol. Cell 65, 25–38 (2017).

5. Hon, C.-C. et al. An atlas of human long non-coding RNAs with accurate 5′ ends. Nature 543, 199–204 (2017).

6. Musacchia, F., Basu, S., Petrosino, G., Salvemini, M. & Sanges, R. Annocript: a flexible pipeline for the annotation of transcriptomes able to identify putative long noncoding RNAs. Bioinformatics 31, 2199–2201 (2015).

7. Lin, M. F., Jungreis, I. & Kellis, M. PhyloCSF: a comparative genomics method to distinguish protein coding and non-coding regions. Bioinformatics 27, i275–i282 (2011).

8. Cabili, M. N. et al. Integrative annotation of human large intergenic noncoding RNAs reveals global properties and specific subclasses. Genes Dev. 25, 1915–1927 (2011).

9. Guttman, M. et al. Chromatin signature reveals over a thousand highly conserved large non-coding RNAs in mammals. Nature 458, 223–227 (2009).

10. Wang, A. & Hai, R. FINET: Fast Inferring NETwork. BMC Res. Notes 13, 521 (2020).

11. Wang, A. & Rong, H. Big-data analysis unearths the general regulatory regime in normal human genome and cancer. bioRxiv 791970 (2019) doi:10.1101/791970.

12. Wang, A. ulncRNA network, https://combai.org/network/lncRNA/.

13. Prescott, S. L. et al. Enhancer divergence and cis-regulatory evolution in the human and chimp neural crest. Cell 163, 68–83 (2015).

14. Rada-Iglesias, A. Is H3K4me1 at enhancers correlative or causative? Nat. Genet. 50, 4–5 (2018).

15. Creyghton, M. P. et al. Histone H3K27ac separates active from poised enhancers and predicts developmental state. Proc. Natl. Acad. Sci. U. S. A. 107, 21931–21936 (2010).

16. Karmodiya, K., Krebs, A. R., Oulad-Abdelghani, M., Kimura, H. & Tora, L. H3K9 and H3K14 acetylation co-occur at many gene regulatory elements, while H3K14ac marks a subset of inactive inducible promoters in mouse embryonic stem cells. BMC Genomics 13, 424 (2012).

17. Liang, G. et al. Distinct localization of histone H3 acetylation and H3-K4 methylation to the transcription start sites in the human genome. Proc. Natl. Acad. Sci. U. S. A. 101, 7357–7362 (2004).

18. Kolasinska-Zwierz, P. et al. Differential chromatin marking of introns and expressed exons by H3K36me3. Nat. Genet. 41, 376–381 (2009).

19. Kouzarides, T. Chromatin Modifications and Their Function. Cell 128, 693–705 (2007).

20. Carlevaro-Fita, J., Rahim, A., Guigó, R., Vardy, L. A. & Johnson, R. Cytoplasmic long noncoding RNAs are frequently bound to and degraded at ribosomes in human cells. RNA 22, 867–882 (2016).

21. Newman, M. E. J. Finding community structure in networks using the eigenvectors of matrices. Phys. Rev. E 74, 036104 (2006).

22. Marquitti, F. M. D., Guimarães, P. R., Pires, M. M. & Bittencourt, L. F. MODULAR: software for the autonomous computation of modularity in large network sets. Ecography 37, 221–224 (2014).

23. Ramilowski, J. et al. Functional Annotation of Human Long Non-Coding RNAs via Molecular Phenotyping. bioRxiv 700864 (2019) doi:10.1101/700864.

24. Parenteau, J. et al. Introns are mediators of cell response to starvation. Nature 565, 612–617 (2019).

25. Morgan, J. T., Fink, G. R. & Bartel, D. P. Excised linear introns regulate growth in yeast. Nature 565, 606–611 (2019).

26. Wei, L.-H. & Guo, J. U. Coding functions of “noncoding” RNAs. Science 367, 1074–1075 (2020).

27. Wang, A. & Rong, H. Noncoding RNAs serve as the deadliest regulators for cancer. Press Cancer Genomics Proteomics RXiv191003934 Q-Bio (2019).

